# No branch left behind: tracking terrestrial biodiversity from a phylogenetic completeness perspective

**DOI:** 10.1101/2022.05.09.491174

**Authors:** Jesús N. Pinto-Ledezma, Sandra Díaz, Benjamin S. Halpern, Colin Khoury, Jeannine Cavender-Bares

**Affiliations:** Department of Ecology, Evolution and Behavior, University of Minnesota, 1479 Gortner Ave, Saint Paul, MN, 55108, USA; Consejo Nacional de Investigaciones Científicas y Técnicas, Instituto Multidisciplinario de Biología Vegetal (IMBIV), CONICET, Universidad Nacional de Córdoba, Córdoba, Argentina; Facultad de Ciencias Exactas, Físicas y Naturales, Universidad Nacional de Córdoba, Córdoba, Argentina; National Center for Ecological Analysis and Synthesis, University of California, Santa Barbara, CA, 93101, USA; Bren School of Environmental Science and Management, University of California, Santa Barbara, CA, 93106, USA; San Diego Botanic Garden, 230 Quail Gardens Dr., Encinitas, CA 92024, USA; International Center for Tropical Agriculture (CIAT), Km 17, Recta Cali-Palmira, Apartado Aéreo 6713, 763537, Cali, Colombia

## Abstract

Biodiversity, as we see it today, ultimately is the outcome of millions of years of evolution; however, biodiversity in its multiple dimensions is changing rapidly due to increasing human domination of Earth. Here, we present the “phylogenetic completeness” (PC) a concept and methodology that intends to safeguard Earth’s evolutionary heritage by maintaining all branches of the tree of life. We performed a global evaluation of the PC approach using data from five major terrestrial clades and compared the results to an approach in which species are conserved or lost randomly. We demonstrate that under PC, for a given number of species extinctions, it is possible to maximize the protection of evolutionary innovations in every clade. The PC approach is flexible and can be used to conduct a phylogenetic audit of biodiversity under different conservation scenarios. The PC approach complements existing conservation efforts and is linked to the post-2020 Convention of Biodiversity targets.

## Introduction

Over more than 3.5 billion years of life on Earth, evolution has generated and honed a vast array of innovations represented by the diversity of genomes and forms across the tree of life. Contemporary species collectively represent the genetic assets that contribute to the functioning of the current biosphere, and these functions in turn serve as the foundation of nature’s contributions to people (NCP) (Díaz *et al*. 2019). Put another way, species embody evolutionary innovations that represent complex and unique approaches to life on Earth. These innovations not only support current NCP, but are also necessary for future benefits to humanity, including those not yet discovered.

Phylogenetic trees depict the hierarchy of life in which species are nested in larger and larger clades, each descended from a more distant common ancestor (Figure 1). They provide information on the breadth and variation of innovations evolution has generated and can be used to inform approaches to species conservation with the goal of minimizing extinction of evolutionary innovations (Faith 2002; Mace *et al*. 2003; Diniz-Filho *et al*. 2013; Larkin *et al*. 2016). Close relatives typically have a high proportion of shared genetics since they arose from a common ancestor at some point in the comparatively recent past, and thus share many of the same innovations.

**Figure 1.**
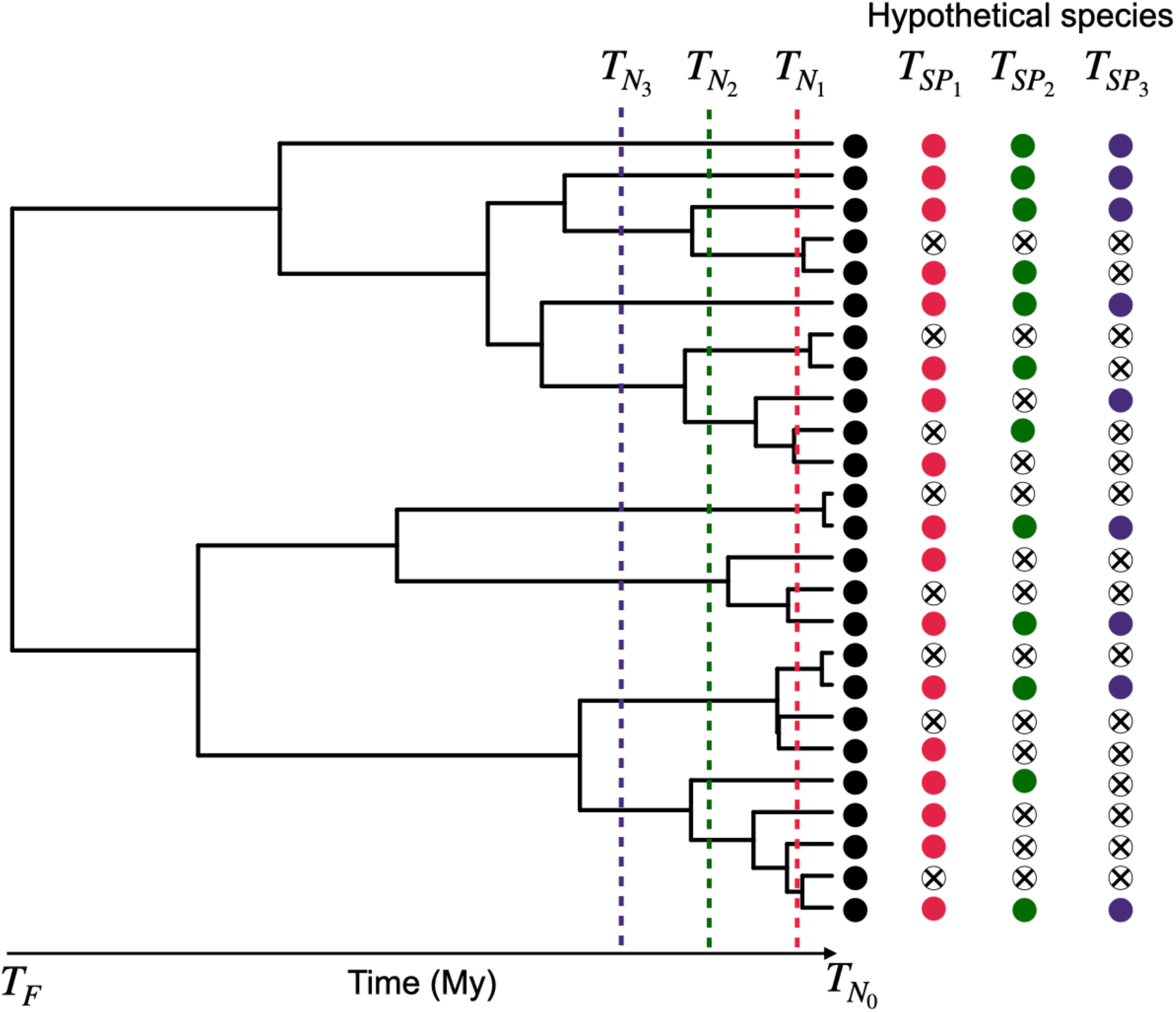
Schematic diagram of the phylogenetic relationships of species within a lineage of organisms, showing the hierarchical nesting of hypothetical species. The number of species that need to be conserved in order to prevent the extinction of any branches in the tree depends on the depth in the phylogeny where we define the branch. Dashed lines represent different ages (T_N_), or depths in the phylogeny, used to define branches—the corresponding number of species preserved (T_SP_), if one species per branch is conserved, is colored the same as the dashed line. For example, compare purple circles (fewer branches and species with a deep slice at T_N3_) to red ones (more branches and species with a more recent slice at T_N1_). The deeper we “slice” the phylogeny, the fewer the species need to be saved in order to preserve all branches of the phylogeny at that slice. Black circles represent species at T_N0_, i.e., the current species, assuming no extinction. Open circles filled with “x” indicate species that went extinct at a specific T_N_.

A wide range of phylogenetic metrics exist that capture diversity across the tree of life (Tucker *et al*. 2017) and are relevant to discerning how much variation is captured under different conservation scenarios. Most evolutionary-based metrics of diversity are applied to community or regional phylogenies and represent different amounts of shared ancestry between taxa (sometimes accounting for other factors, such as probability of extinction). The canonical example is phylogenetic diversity (PD, *sensu* (Faith 1992)), which, for a subset of species, is defined as a sum of all branch lengths required to connect those species or the total amount of evolutionary history contained by those species (Faith 1992, 2002). Since its introduction in 1992, PD has been used in ecological and conservation studies as a measure of biodiversity that captures information about the number of accumulated features in the branches of the tree of life (Scherson and Faith 2018).

In our era of rapid biodiversity loss (Tilman *et al*. 2017; IPBES 2019; Díaz *et al*. 2019) keeping all species—the tips of the tree of life—is not realistic, and indeed is practically unfeasible given recent extinction rates (Crozier 1997; Rounsevell *et al*. 2020). However, it may still be possible to conserve most of the branches of the tree of life, assuming it is possible to meaningfully define those branches (Figure 1). This concept is the principle behind phylogenetic completeness, a concept and methodology we propose here that aims to preserve Earth’s evolutionary heritage (*sensu* Mooers *et al*. 2005) by maintaining all branches across the tree of life. The approach aims to facilitate a phylogenetic audit of biodiversity. The central goal is to track all of life’s variation that has arisen over the course of evolution, and still remains, in both early-diverged lineages and recently-derived lineages in each of the major living clades and across the entire tree of life. Phylogenetic completeness (PC) and phylogenetic diversity (PD) both aim to maintain as many branches as possible from the tree of life, recognizing it is not possible to retain all species. But given the hierarchical nature of phylogenetic trees, how a lineage is circumscribed—or what constitutes a “branch” that should be protected—can influence the outcome (Vane-Wright *et al*. 1991; Diniz-Filho *et al*. 2013; Scherson and Faith 2018). PC differs from PD in that it slices across the tree at a given point in evolutionary history and defines a set of branches based on this cut off (Figure 1), using a phylogenetic accounting framework to identify and audit all distinctive branches in the tree of life at local, regional or global scales, rather than only maximizing phylogenetic breadth.

Here we conducted a series of analyses to explore the implications for conservation depending on the depth in the tree of life where a “branch” is defined. The goal was to develop and apply a framework for conservation of species that minimizes hemorrhaging of Earth’s evolutionary assets, given a fixed level of species loss that is assumed to be unavoidable. We then compared the loss of PD under conservation scenarios in which the set of species targeted for conservation was based on an “informed” phylogenetic completeness approach or in which species conserved or lost to extinction occurred randomly.

## Methods

### Phylogenetic completeness (PC) approach

We developed an approach in which a phylogeny is iteratively sliced at different periods of time (T_N_) until a specified finish time (T_F_). For example, if a phylogeny is sliced every T_N_ = 2 million years until T_F_ = 50 million years, a total of 25 slice points is obtained (see also Figure 1). These slice points are then used to drop all but one terminal tips—or operational taxonomic units (OTU)—from the phylogeny; with this approach, we ensure that at least one descendant OTU of each lineage at a specific time (T_N_) is kept. In other words, by keeping at least one OTU from each lineage in the tree of life, we aim to maximize the preservation of the deepest evolutionary history. At each slice point (T_N_) we additionally calculated the number of species (T_SP_) and phylogenetic diversity (T_PD_) as the simple sum of branch lengths at the specific slice point (T_N_).

### Empirical assessment

The empirical assessment focused on five major terrestrial clades (seed plants, amphibians, squamates, birds, and mammals). Description of the data and software used can be found in the WebPanel 1.

We tested the reliability of the PC approach by slicing each phylogeny every T_N_ = 100 Ky until T_F_ = 100 My and calculated the T_SP_ and T_PD_ at every slice point. These metrics were then used to identify change points in the phylogenetic diversity over the 100 My time period. Change points were evaluated using Bayesian Multiple Changing Points (MCP) regressions. The first changing point plus its credible intervals (CIs) identified by the MCP analysis were used as cutoff thresholds to estimate the number and identities of OTUs to be kept. This procedure allowed us to identify different change points or cutoff thresholds in the phylogenetic diversity over 100 My for each clade separately and consequently prevent us establishing a fixed arbitrary cutoff threshold (e.g., setting a changing point at 2 My as cutoff threshold) for all clades.

We compared the diversity in each clade for a PC conservation scenario in which species were managed to maintain all phylogenetic branches to random losses (RAND). In other words, we removed OTUs at random until the identified cutoff threshold for each clade separately. This procedure was repeated 1000 times, and at each step the T_SP_ and T_PD_ were estimated.

Finally, using the OTUs identities from both the PC and the random loss scenarios we mapped the phylogenetic diversity of seed plants and terrestrial vertebrates globally. These maps were used to estimate the difference (Δ*PD*) between the observed PD (PD_OBS_) and the expected PD (PD_EXP_) under either the phylogenetic completeness (PD_PC_) and the random loss approach (PD_RAND_).

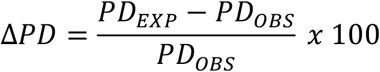

These maps represent the proportional difference between the observed (PD_OBS_) and expected (PD_PC_ or PD_RAND_) phylogenetic diversity, where negative values suggest that a grid cell will lose a proportion of its PD according to a specific conservation scenario, e.g., under the PC scenario. Note that the lower CI from the Bayesian MCP regressions were used as variable cutoff thresholds for each clade for mapping purposes.

## Results

### Phylogenetic completeness

Bayesian MCP models revealed variable cutoff thresholds for each clade (WebFigure 1; WebTable 1). Based on these thresholds, losses of species ranging from 1.34% to 18.11% are estimated to occur in each of the major clades (seed plants, amphibians, squamates, birds, and mammals) (WebTable 1) while still safeguarding between 97.27% to 99.97% of the phylogenetic diversity—i.e., of the evolutionary history of each clade (WebTable 1; Figure 2). If the lower credible interval (LCI or the 2.5% quantile of the posterior distribution) of our Bayesian model estimates were used to define the phylogenetic branches to be conserved (Figure 2; WebTable 1), a higher number of species and branches in the tree of life would be safeguarded (Table 1).

**Figure 2.**
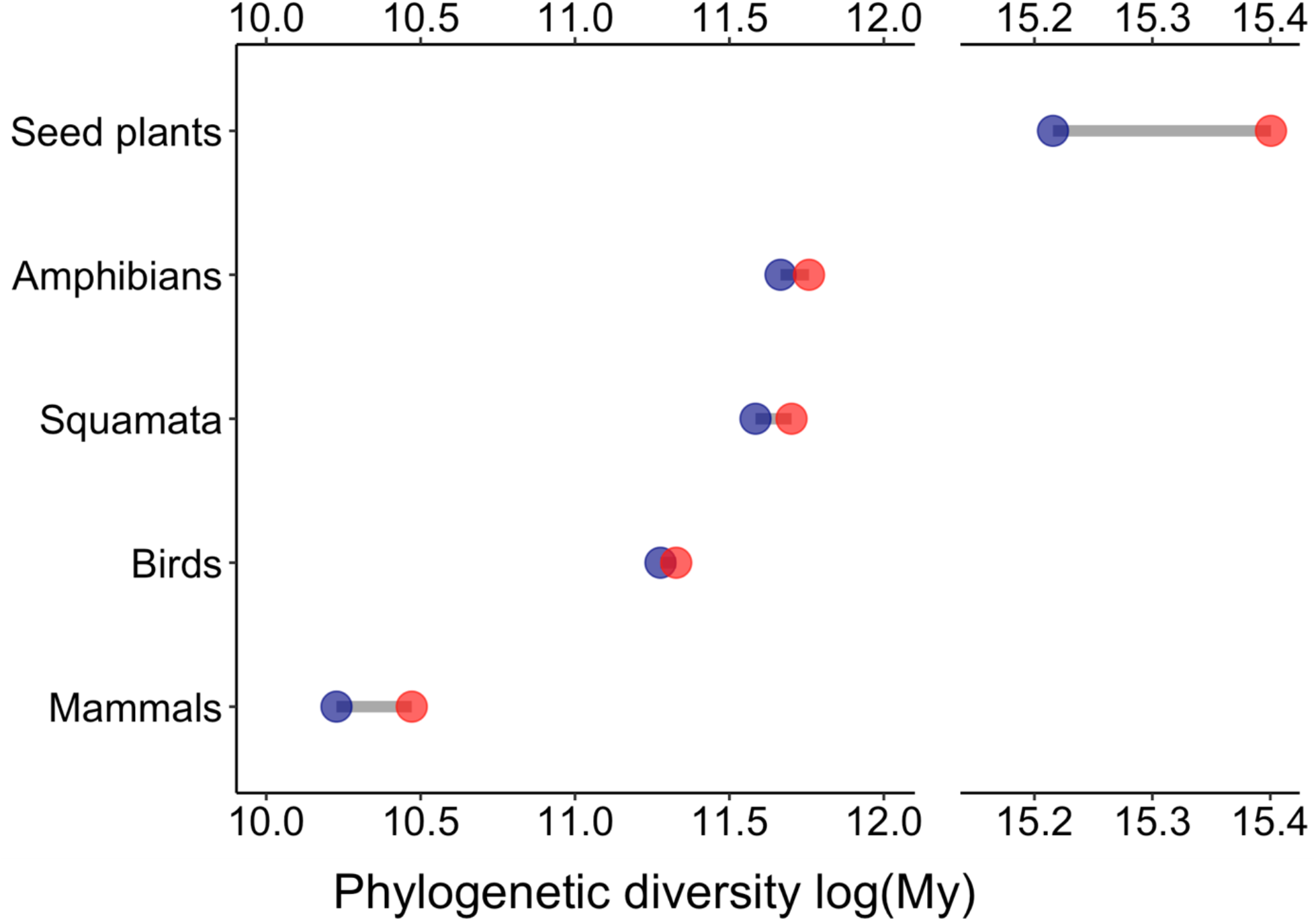
Difference between the remaining phylogenetic diversity under phylogenetic completeness (red) and random loss (blue) scenarios for five major terrestrial clades (seed plants, amphibians, squamates, birds, and mammals). The X-axis was log-transformed for plotting purposes. In all cases, the phylogenetic diversity is higher for the phylogenetic completeness approach.

**Table 1.** Number of branches (NB) conserved under the scenarios of phylogenetic completeness (PC) and random loss (Rand) conservation scenarios. The number of branches conserved under the PC approach is higher than the random loss scenario for the same number of species except for birds and mammals. Percentages of NB under PC and Rand scenarios are displayed within brackets. Number of branches within a clade assuming no extinction is displayed for reference. The threshold number of species under either the PC or Rand scenarios corresponds to the number of species at the lower CI identified using the Bayesian MCP regressions.

| Clade | Observed N species | Threshold N species | Threshold (My) | NB all species | NB-PC | NB-Rand |
| --- | --- | --- | --- | --- | --- | --- |
| Seed plants | 353185 | 312540 | 1.76 | 438863 | 397974 [90.68%] | 391851 [89.29%] |
| Amphibians | 7238 | 5927 | 4.43 | 14474 | 12614 [87.15%] | 11852 [81.88%] |
| Squamata | 9755 | 8141 | 2.61 | 19508 | 17186 [88.10%] | 16280 [83.45%] |
| Birds | 9993 | 9859 | 0.33 | 19984 | 19716 [98.66%] | 19716 [98.66%] |
| Mammals | 5911 | 5802 | 0.31 | 11820 | 11602 [98.16%] | 11602 [98.16%] |

Note: the observed number of species (second column) corresponds to the number of species sampled in each phylogenetic tree and might not represent each clade's true number of species.

These analyses demonstrate that if conservation efforts are focused on maintaining defined branches of the tree of life it is possible to maximize the accumulated evolutionary innovations that are safeguarded across all clades even when individual species go extinct. Figure 2 shows the comparison between the estimates of PD under both phylogenetic completeness (PC) and random loss (RAND) scenarios. Under PC, a higher number of branches (Table 1) and greater evolutionary history in each clade is preserved for a given number of species extinctions (Figure 2; WebFigure 2).

Spatial patterns of ΔPD under PC and RAND scenarios (Figure 3; WebFigure 3), show how conservation informed by PC safeguards a greater proportion of evolutionary history even with the same number of species extinctions (WebFigure 2). For example, for seed plants in tropical regions across the world, conservation informed by PC resulted in PD loss below 10%, whereas the Rand scenario resulted in 10-20 % of PD loss. Extinction patterns of terrestrial biodiversity under PC and Rand scenarios at the biome level (WebFigure 4) also show greater preservation of accumulated evolutionary innovations when conservation is targeted to maintain branches of the tree of life. Nevertheless, we find that Tundra and Taiga biomes are susceptible to high losses in PD, especially for seed plants, amphibians, and mammals, even under PC scenarios. Indeed, further statistical analyses revealed small to no evidence in favor of the phylogenetic completeness over random loss (evidence ratio < 5) in these biomes (WebTable 2). In contrast, tropical biomes (for both forest and grasslands) show limited losses in PD for the same threshold values used to define branches, as in Tundra and Taiga biomes (WebFigure 4; WebTable 2). These results indicate that for a given number of species extinctions, tropical biomes will lose fewer branches of the tree of life and are thus less susceptible to loss of evolutionary history.

**Figure 3.**
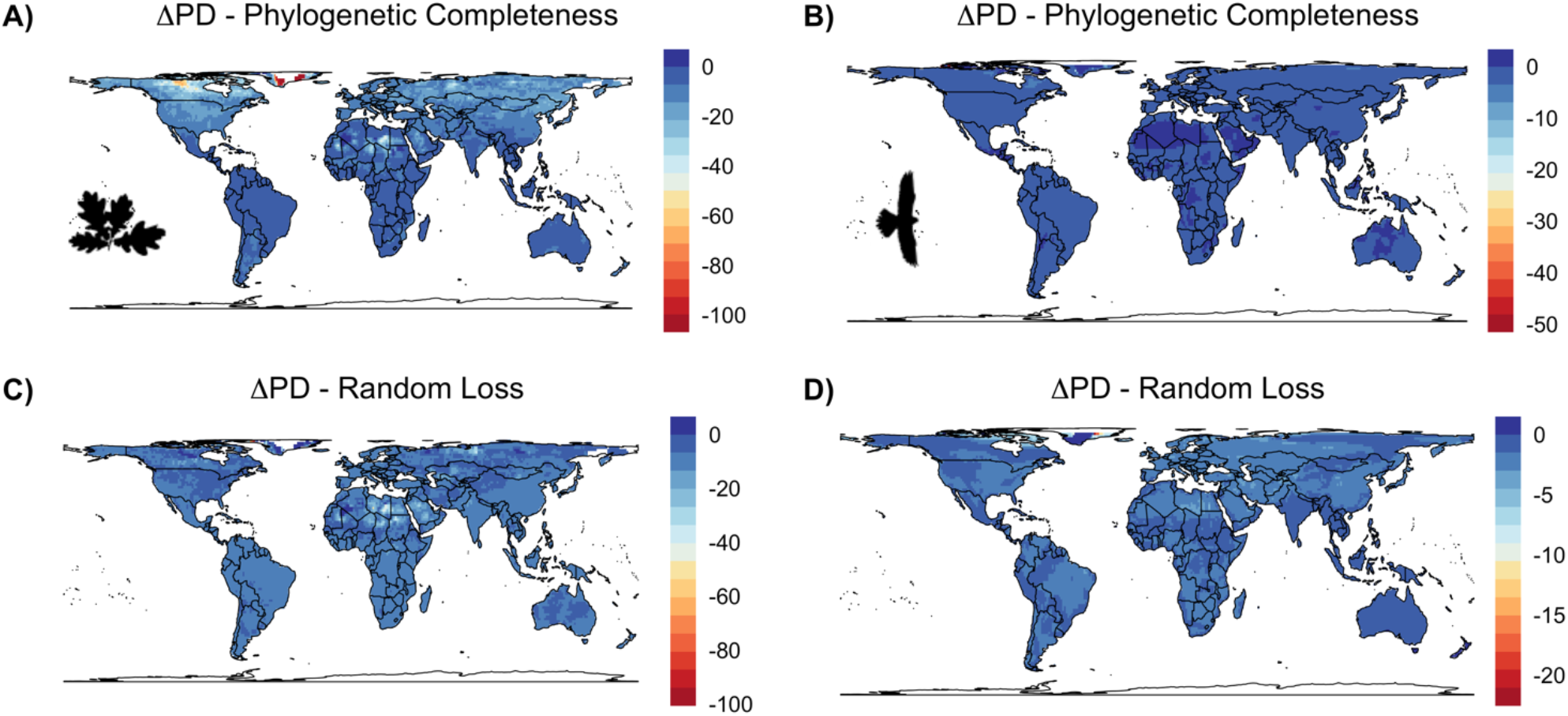
Mapped phylogenetic diversity for seed plants (A-C) and birds (B-D) globally under scenarios of phylogenetic completeness and random loss. Legends indicate the proportional loss of phylogenetic diversity (Δ*PD*). Blue tones indicate that more branches of the tree of life have been preserved and red tones that more branches have been lost. Comparing the two scenarios, globally more branches of the tree of life are conserved in the phylogenetic completeness scenario for the same number of species extinctions. Maps for amphibians, squamates, and mammals can be found in the supplementary material (WebFigure 3).

## Discussion

Our “phylogenetic completeness” framework for informing biodiversity conservation focuses on maintaining the accumulated evolutionary innovations across the tree of life with the intent of leaving no branch behind. We introduce a rigorous approach for defining branches across clades of terrestrial organisms to ascertain where in the tree of life there is high evolutionary redundancy and where a single species may represent an entire branch. In doing so, we establish phylogenetic branches as units of conservation priority rather than specific species or OTUs. By defining these branches and the species contained within them, the phylogenetic completeness approach provides critical information on branches at risk of extinction where there is low redundancy as well as flexibility in which species could be targeted for conservation in cases of high redundancy (Mooers *et al*. 2005).

The approach is particularly useful in developing priorities for *ex situ*, as well as for *in situ* in protected areas, by tracking which branches of the tree of life are currently safeguarded and by identifying the branches that are at highest risk—those not currently protected or with the least amount of their range protected. The Kunming-Montreal Global Biodiversity Framework, just adopted by the Convention on Biological Diversity aims to reduce the extinction rate and risk of all species by tenfold by 2050 (Goal A) (CBD 2022), but does not explicitly include the tree of life. Yet calls by the international scientific community to set ambitious goals for biodiversity recognize that not all extinctions have equal consequences and that phylogenetic dimensions of biodiversity need to be recognized among the criteria for implementation (Díaz *et al*. 2020). The PC approach is consistent with the overall the CBD agenda because, at a given overall extinction rate, it can help identify what species or clades will represent disproportionally large or small losses of evolutionary history. As well as having self-evident intrinsic value, evolutionary history is considered a major basis of “maintenance of options”, one of the major categories of nature’s contributions to people recognized by the intergovernmental biodiversity science-policy interface (IPBES 2019).

Increasing human domination of Earth and its ecosystems is rapidly changing biodiversity patterns and negatively impacting the biosphere’s capacity to provide essential contributions to humanity (Tilman *et al*. 2017; Díaz *et al*. 2019). Safeguarding all remaining biodiversity, although desirable, is unrealistic on the basis of virtually all projections (Pimm *et al*. 2014; Tilman *et al*. 2017; Scherson and Faith 2018); the footprint of humanity is currently too large to completely avoid further extinctions. Scientists have recognized the challenge of developing logical conservation solutions given the complexity of stakeholders, managers, and indirect actors as a ‘wicked problem’ that has no straightforward solution (Vane-Wright *et al*. 1991; Crozier 1997; Mace 2014). Focusing on the conservation of evolutionary history has been hailed as an integrative way to safeguard most of the biodiversity and its functions (Faith 1992; Mooers 2007; Mishler *et al*. 2014; Larkin *et al*. 2016). For example, a recent study by Molina-Venegas and collaborators (Molina-Venegas *et al*. 2021) (Molina-Venegas *et al*. 2021) found strong evidence that plant evolutionary history is tightly linked to multiple plant use categories and therefore to human well-being (Molina-Venegas *et al*. 2021). These findings, among others, support the idea that conserving evolutionary history is critical for future human well-being (Forest *et al*. 2007; Molina-Venegas *et al*. 2021).

We demonstrate that under the PC framework, the loss of evolutionary history— measured as the total branch lengths in the tree of life—can be minimized (Figure 2, 3). Our results show that the loss of evolutionary heritage for five major terrestrial clades is lower compared to phylogenetically random extinctions (Figure 2, WebFigure 2). In addition, although current species extinctions are non-random (Purvis *et al*. 2000), evidence suggests that the impact of random extinctions erases more evolutionary history on clades with imbalanced phylogenetic trees (Heard and Mooers 2002) as the ones assessed in this study.

By assessing the evolutionary history loss at biome level (WebFigure 4) we found that PC does not outperform the conservation scenario of random extinction in biomes at high latitudes (e.g., Tundra and Taiga), in particular for the seed plants, amphibians, and squamates (WebTable 2). At the scale of the biome, the loss of evolutionary history under the PC framework seems to have acted selectively—extinctions are concentrated in specific taxa— causing those depauperate lineages in these biomes to become even more depauperate. Indeed, at high latitudes an individual species frequently represents an entire phylogenetic branch while at low latitudes (tropics), a branch is likely to contain many species. This pattern is largely the consequence of more recent divergence times and higher speciation rates in the tropics.

However, the spatial scale (grain size) must also be considered; for example, one hectare in a tropical forest can hold ∼650 tree species, which is more than all tree species that occur at high latitudes (Coley and Kursar 2014). Despite this high diversity, tropical forests are usually hyper-dominated by a fraction of species (∼1.4% of about 16,000 tree species estimated for the Amazonian Forest are considered as hyper-dominant) that are specialists to their habitats and have large geographical ranges (ter Steege *et al*. 2013). The less abundant species or poorly known species with small geographical ranges are potentially threatened.

Although it is beyond the scope of this article to conduct, PC audits at local and regional scales could help to identify which species may be prioritized to prevent losing branches of the tree of life at local and regional scales. To demonstrate scaling our approach to local sites, we assessed a plant community-level phylogeny from the Cedar Creek Ecosystem Science Reserve (Pinto-Ledezma *et al*. 2020). This analysis reveals which taxa are unique and which taxa present evolutionary redundancy, i.e., red branches in WebFigure 5. Moreover, using plant community composition (N = 987 plots of 40 × 40 m) from the National Ecological Observatory Network (NEON) we found that under scenarios where extinctions are predicted to occur all branches are preserved according to our phylogenetic accounting framework. Our results demonstrate that phylogenetic diversity is nearly equivalent to that calculated from observed data where no extinctions are assumed to have occurred (WebFigure 6). This additional analysis reveals that, despite variation in the number of species lost (WebFigure 6A), focusing on branches as units of conservation aids in maintaining the evolutionary history of local communities.

Multiple approaches have been proposed to assess changes in biodiversity focusing on “hotspot” areas (spatial prioritization), or taxa (taxonomic prioritization) for conservation purposes (Margules and Sarkar 2007). These approaches rely on the use of metrics that capture different dimensions of biodiversity, e.g., metrics that capture evolutionary changes among a set of taxa (Rodrigues *et al*. 2005; Margules and Sarkar 2007) or the variation in form and function of taxa within communities (Díaz and Cabido 2001; Petchey and Gaston 2006). Despite their usefulness for assessing the state and the fate of biodiversity, most of these metrics, if not all, are sensitive to information (in)completeness. Missing information can result in misleading metric calculations and inappropriate interpretations of spatial or taxonomic comparisons (Isaac 2004; Rodrigues *et al*. 2011; Diniz-Filho *et al*. 2013; Weedop *et al*. 2019). The PC framework introduced here represents a complementary approach to counting numbers of species or comparing levels of phylogenetic diversity to assess biodiversity under alternative conservation scenarios. It provides an accounting framework that prioritizes conservation of branches of the tree of life rather than individual taxa (Table 1; Figure 3). It also allows the identification of areas susceptible to high losses of evolutionary heritage (Figure 3; WebFigure 3), which can be used as baseline information for spatial prioritization, providing a broader context for local decision making (Mace *et al*. 2003; Mishler *et al*. 2014; Chaplin-Kramer *et al*. 2022; Silvestro *et al*. 2022).

Moreover, given that the PC framework focuses on the branches of the tree of life, in the context of spatial prioritization, our approach my help existing spatial phylogenetic approaches—e.g., categorical analysis of neo- and paleo-endemism (Mishler *et al*. 2014; Thornhill *et al*. 2016), phylogenetic endemism (Rosauer *et al*. 2009)—to deal with the issue of missing data. To illustrate this point, if we consider protecting at least one descendant taxon from a specific node in the phylogeny, this taxon contains information—genetic, traits, functions— that captures most of the evolutionary history of the lineage, i.e., the evolutionary history of the taxon plus its common ancestor (see Figure 1; WebFigure 5). If species within the branch have not yet been identified or are not readily observed, the branch itself is still preserved, with the caveat that phylogenetic information remains imperfect—i.e., Darwinian shortfall in biodiversity and conservation (Diniz-Filho *et al*. 2013). By focusing on branches of the tree of life as an additional conservation criterion beyond those prioritized in other ways, it is possible to reduce the impact of missing data and taxonomic inflation on evolutionary diversity estimations and conservation planning (Isaac 2004; Rodrigues *et al*. 2011; Diniz-Filho *et al*. 2013; Allen and Mishler 2022).

Finally, we are aware that our empirical evaluation was based on incomplete data, but it is the best phylogenetic and geographical data available to date. Although our estimations are robust, more research may be needed to fully understand the potential of the PC framework in biodiversity conservation. In addition, this study is limited to terrestrial biodiversity; thus, evaluations of aquatic biodiversity are required. Future studies focusing on aquatic systems or terrestrial biodiversity not considered here (e.g., Hexapoda) may help to refine the PC’s potential in biodiversity assessment and conservation. These studies should also investigate the relative impact of branch selection, i.e., “the agony of choice,” or which branch should be prioritized and its conservation implications. In other words, where should we focus our efforts to minimize the loss of evolutionary heritage. Lastly, we emphasize that phylogenetic completeness is not a metric of phylogenetic diversity but an accounting framework that identifies targets (unique branches) for eco-evolutionary or conservation questions in a phylogenetic context. Once the unique branches are identified, existing metrics can be used (WebFigure 6) to estimate the impact of lineage loss on biodiversity estimations, conservation scenarios or ecological research.

### Moving forward

Phylogenetic completeness provides a means to account for phylogenetic branches and facilitates the safeguarding of evolutionary heritage, adding to but not replacing other priorities for conservation. For example, the approach will highlight when a single species is the lone member of a branch, increasing its conservation priority. The spatial scale of evaluation is important to consider, and application at local scales in coordination with regional scales is critical. It would be dangerous to consider that maintaining only one species in a branch is adequate at large scales, mainly because some species were extirpated locally and currently occur in a fraction of their original geographic distribution (e.g., Hyacinth macaw, African lion). However, an informed and coordinated conservation may allow individual protected areas to focus on conservation of species that maintain all branches locally, emphasizing particular species within a branch based on other conservation criteria. An auditing procedure with appropriate tools would enable coordination at regional scales to ensure that most or all species in a particular branch are being managed. By focusing on branches of the tree of life as an additional conservation criterion beyond those prioritized in other ways, we will be less likely to overlook the importance of management for taxa that alone represent an evolutionary branch (Mace *et al*. 2003; Mace 2014; Allen and Mishler 2022).

## Conclusion

In our era of rapid global change and rapid biodiversity loss, conservation efforts must reckon with the reality that we will not succeed in saving all species on Earth. We outline an approach for conservation, which we call phylogenetic completeness, that focuses on saving the accumulated innovations that have evolved in Earth’s biota by counting individual branches in the tree of life as units of conservation priority. The approach benefits from detailed information of the tree of life that is only now sufficiently resolved to be applicable to all of life on Earth.

The approach complements other conservation efforts and is directly relevant to the targets of the Kunming-Montreal Global Biodiversity Framework established by the Convention on Biological Diversity.

## Acknowledgements

Support to this project was provided by the National Science Foundation (NSF) through the Macrosystems Biology & NEON-Enabled Science program (DEB-2017843). Further support was provided by the NSF BII ASCEND (DBI-2021898). The work is part of the FITBITs working group hosted by the National Center for Ecological Analysis and Synthesis.

## Statement of Authorship

All authors contributed intellectually to the manuscript. J.C.-B. and J.N.P.-L. conceived of and framed the manuscript. J.N.P.-L. performed all the statistical and spatial analyses with input from J.C.-B. All authors edited the manuscript.

## Open Research

### Data Availability Statement

All data used in this manuscript are publicly available. Main sources are provided in the WebPanel 1. R functions and examples for data analyses are publicly available at https://github.com/jesusNPL/FITBITs.

