## Supplemental Table 1 for "No branch left behind: tracking terrestrial biodiversity from a phylogenetic completeness perspective": Pinto-Ledezma_EtAl_SuppInfo_WebFigure_S1.docx

**JN Pinto-Ledezma *et al*. – Supporting Information**

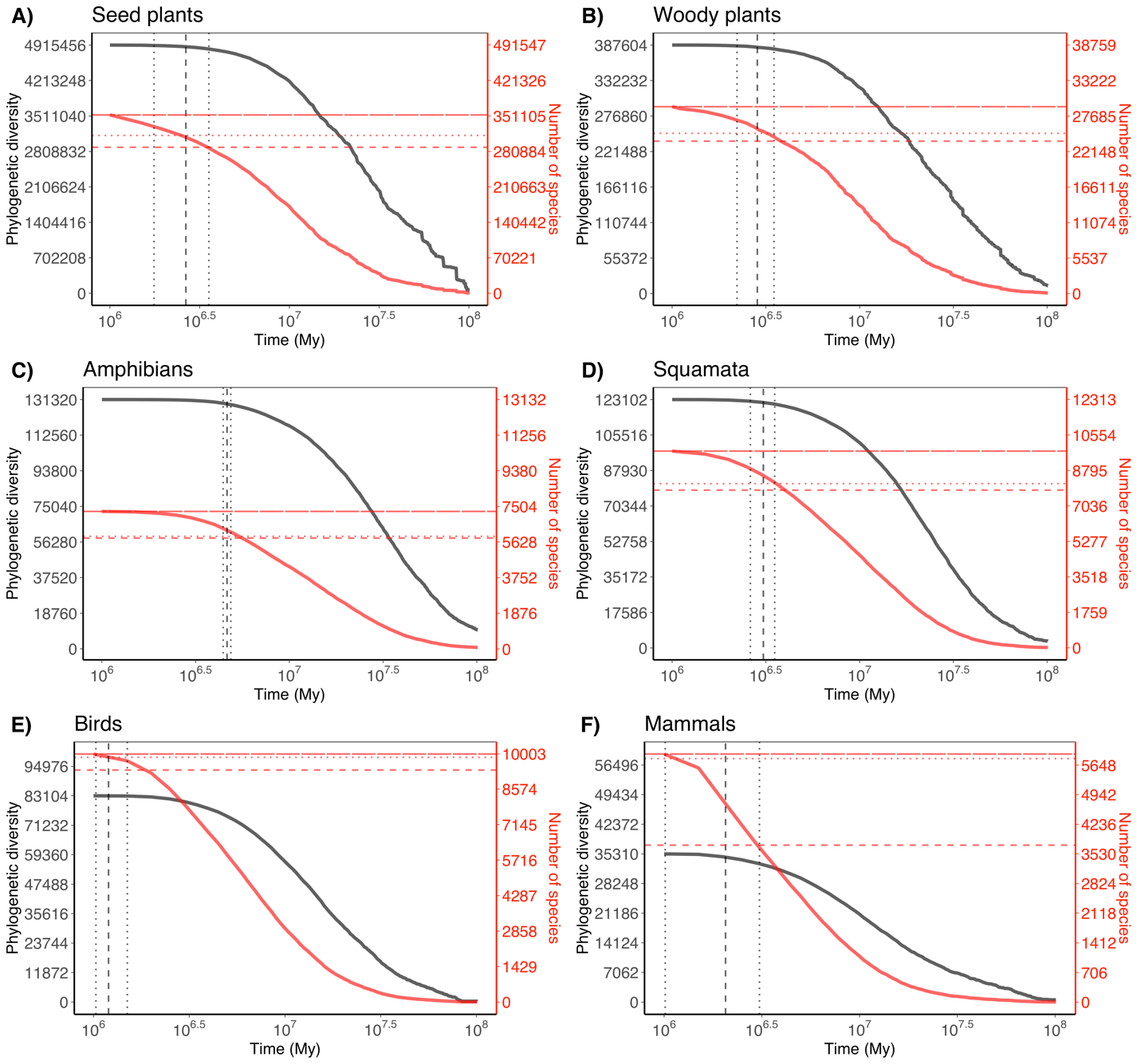

**WebFigure 1**. Profiles of phylogenetic diversity and number of species over 100 million years for terrestrial biodiversity, including seed plants (A), woody plants (B), amphibians (C), squamates (D), birds (E), and mammals (F). Black dashed vertical lines represent the first changing point identified by the Bayesian Multiple Changing Points (MCP) regressions. Black dotted vertical lines represent the 95% credible intervals. Red long-dashed horizontal lines represent the observed number of species for each clade. Red dashed horizontal lines are the number of species expected under the first changing point. Red dotted horizontal lines represent the expected number of species under the lower bound (or 2.5% credible interval) of the first changing point. See WebTable 1 for a comprehensive numerical summary of the changing points.
