## Supplemental Table 1 for "No branch left behind: tracking terrestrial biodiversity from a phylogenetic completeness perspective": Pinto-Ledezma_EtAl_SuppInfo_WebFigure_S2.docx

**JN Pinto-Ledezma *et al*. – Supporting Information**

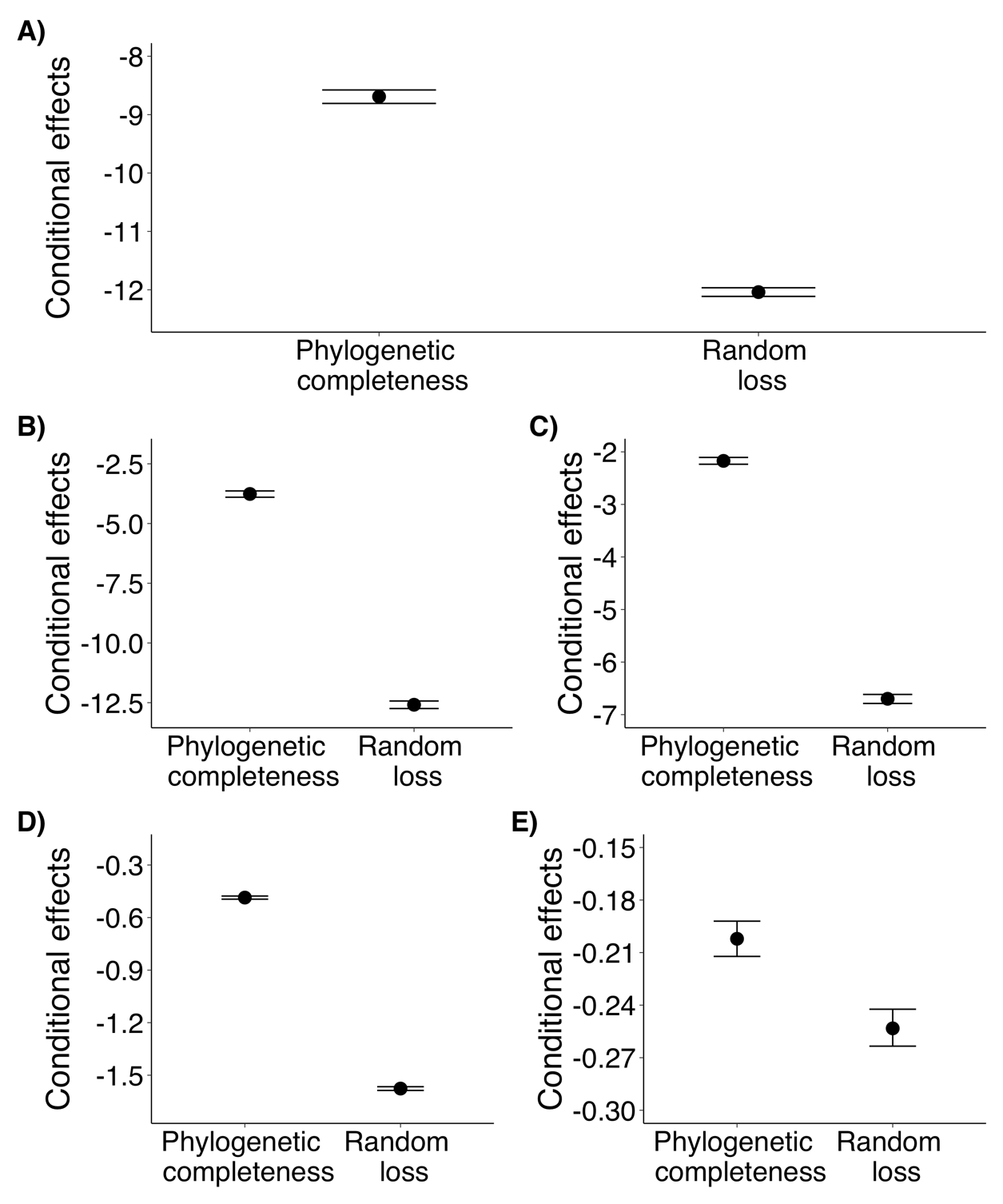

**WebFigure 2**. Statistical comparison between phylogenetic completeness and random loss. In all cases, phylogenetically random extinctions cause a higher loss of evolutionary history. (A) seed plants, (B) amphibians, (C) squamates, (D) birds, and (E) mammals.
