## Supplemental Table 1 for "No branch left behind: tracking terrestrial biodiversity from a phylogenetic completeness perspective": Pinto-Ledezma_EtAl_SuppInfo_WebFigure_S3.docx

**JN Pinto-Ledezma *et al*. – Supporting Information**

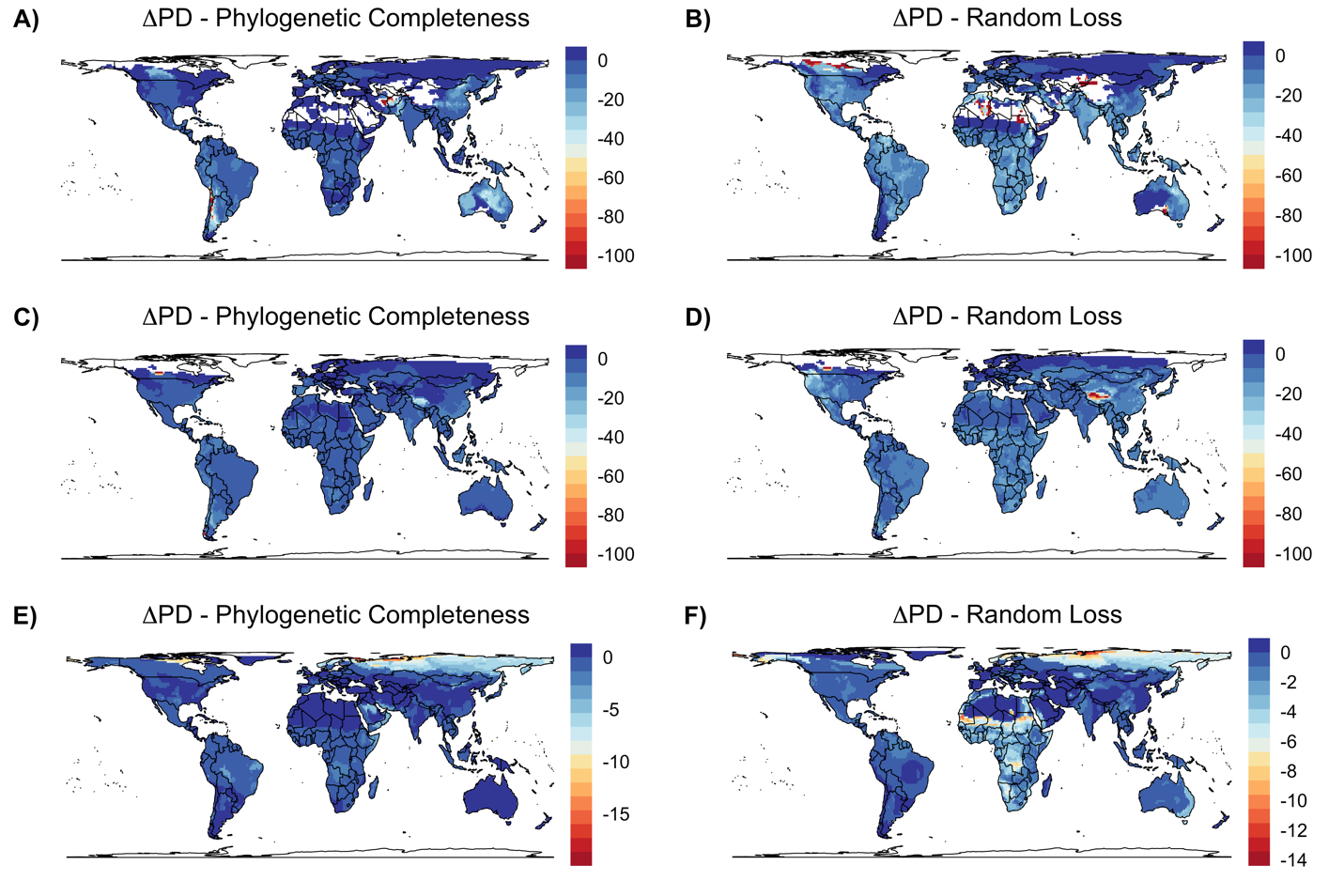

**WebFigure 3**. Mapped phylogenetic diversity under scenarios of phylogenetic completeness and random loss for amphibians (A-B), and squamates (C-D), and mammals (E-F). Legends indicate the proportional loss of phylogenetic diversity ($\Delta PD$). Blue tones indicate that more branches of the tree of life have been preserved and red tones that more branches have been lost.
