## Supplemental Table 1 for "No branch left behind: tracking terrestrial biodiversity from a phylogenetic completeness perspective": Pinto-Ledezma_EtAl_SuppInfo_WebFigure_S4.docx

**JN Pinto-Ledezma *et al*. – Supporting Information**

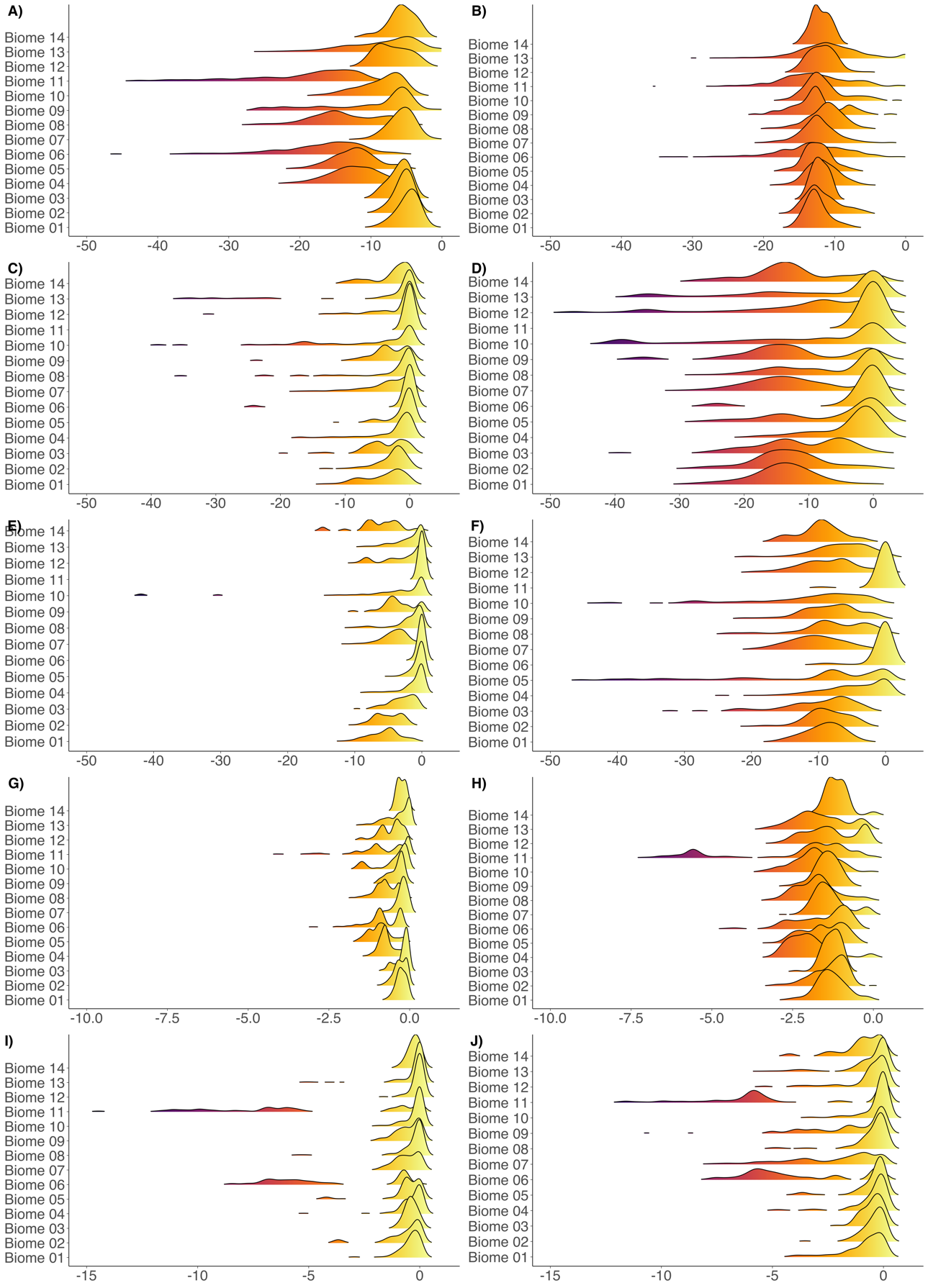

**WebFigure 4**. Patterns of ΔPD at biome level. Ridge gradients represent the pixel values within biomes for vascular plants in the Americas and the terrestrial vertebrates globally. Yellow tones indicate no difference between the observed PD and the expected PD under the phylogenetic completeness or random loss. Red tones suggest higher difference or high proportion of phylogenetic diversity loss. Biome codes: Biome 01 = “Tropical & Subtropical Moist Broadleaf Forests”; Biome 02 = “Tropical & Subtropical Dry Broadleaf Forests”; Biome 03 = “Tropical & Subtropical Coniferous Forests”; Biome 04 = “Temperate Broadleaf & Mixed Forests”; Biome 05 = “Temperate Conifer Forests”; Biome 06 = “Boreal Forests/Taiga; Biome 07 = “Tropical & Subtropical Grasslands, Savannas & Shrublands”; Biome 08 = “Temperate Grasslands, Savannas & Shrublands”; Biome 09 = “Flooded Grasslands & Savannas”; Biome 10 = “Montane Grasslands & Shrublands”; Biome 11 = “Tundra”; Biome 12 = “Mediterranean Forests, Woodlands & Scrub”; Biome 13 = “Deserts & Xeric Shrublands”; and Biome 14 = “Mangroves”.
