## Supplemental Table 1 for "No branch left behind: tracking terrestrial biodiversity from a phylogenetic completeness perspective": Pinto-Ledezma_EtAl_SuppInfo_WebFigure_S5.docx

**JN Pinto-Ledezma *et al*. – Supporting Information**

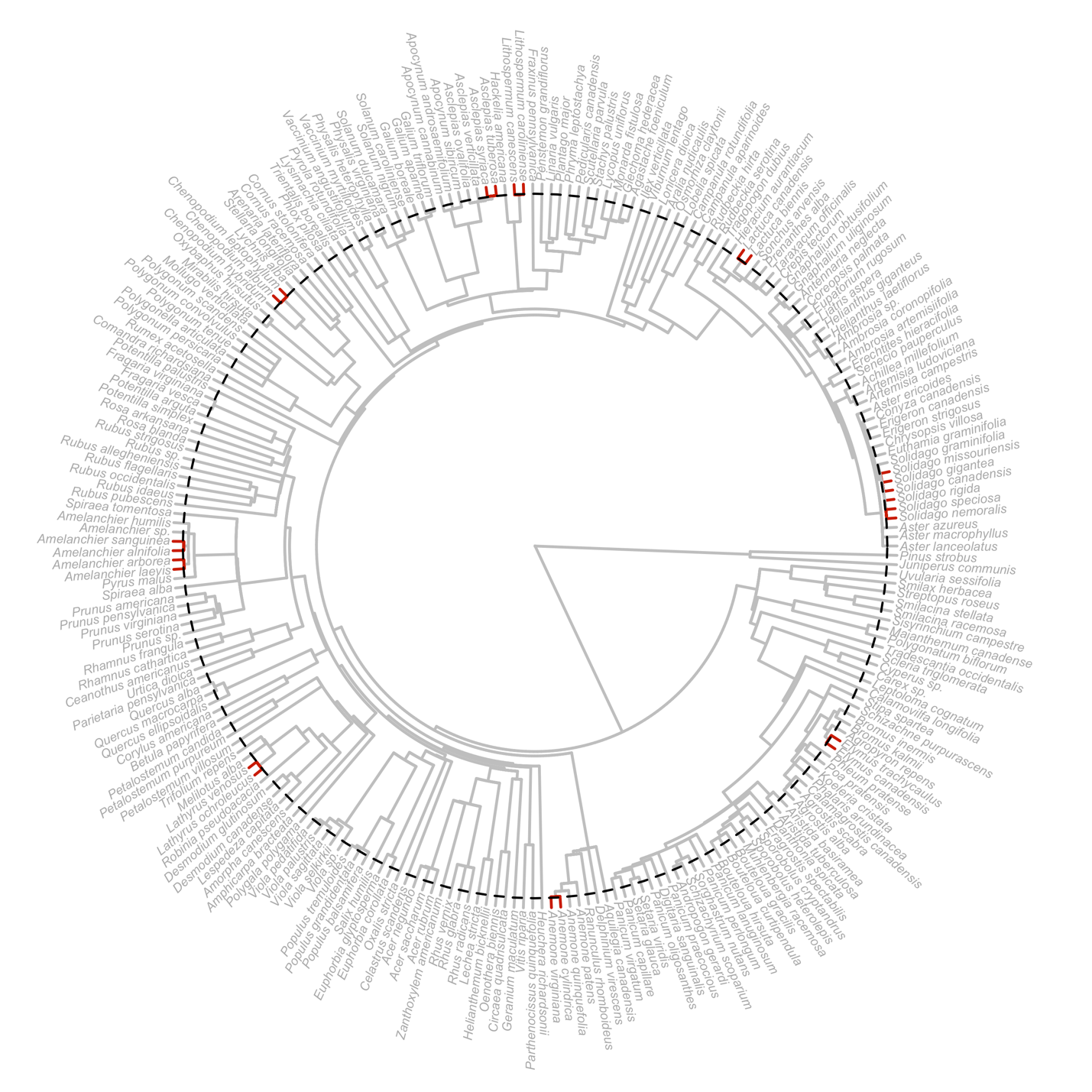

**WebFigure 5.** Community-level phylogenetic hypothesis for seed plant species (N = 243) at the Cedar Creek Ecosystem Reserve. Red-colored terminal branches indicate the taxa that present evolutionary redundancy. The black dotted-line represents the threshold of 1.76 My identified for seed plants.
