## Supplemental Table 1 for "No branch left behind: tracking terrestrial biodiversity from a phylogenetic completeness perspective": Pinto-Ledezma_EtAl_SuppInfo_WebFigure_S6.docx

**JN Pinto-Ledezma *et al*. – Supporting Information**

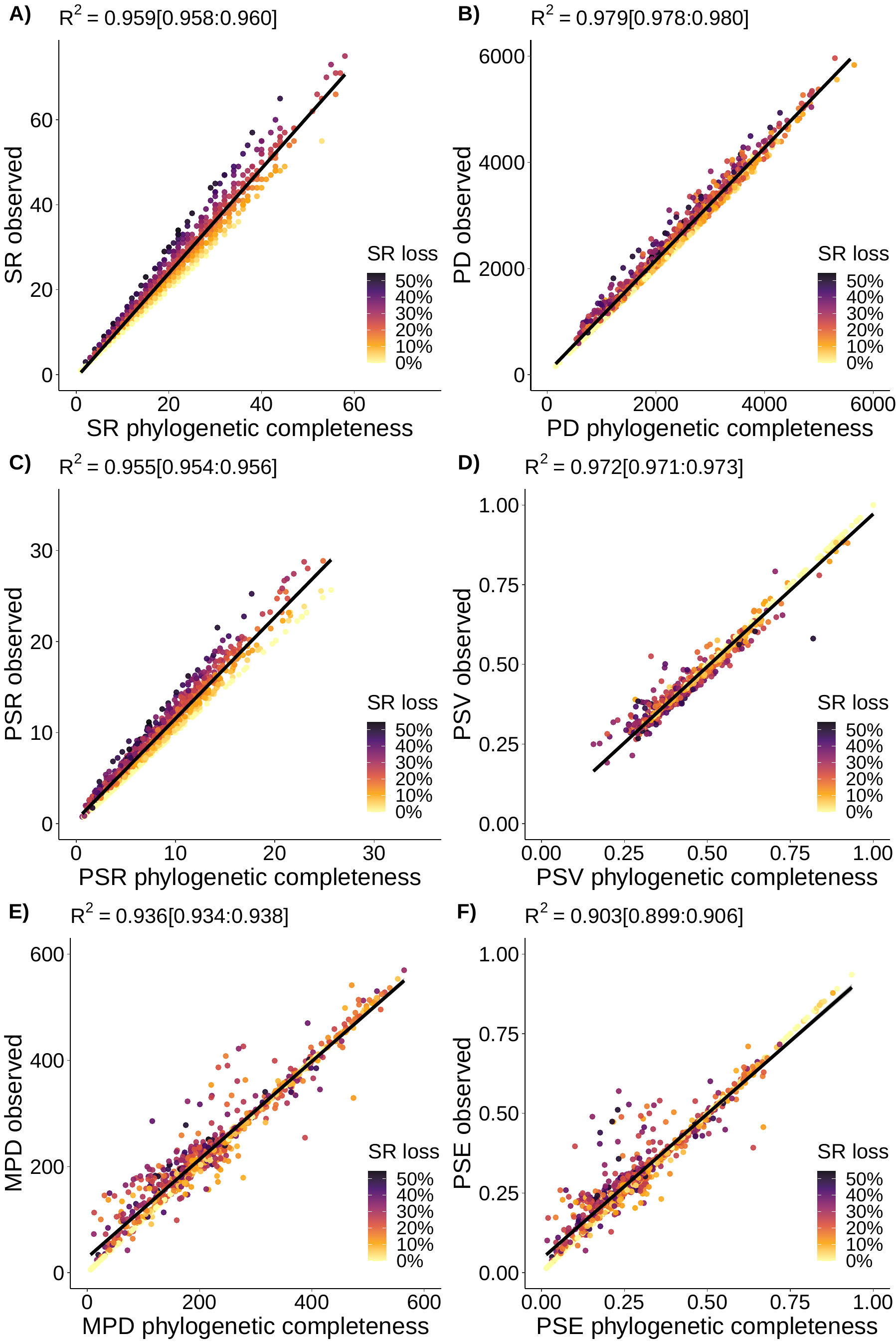

**WebFigure 6.** Association between metrics of phylogenetic diversity under phylogenetic completeness (x-axes) and metrics of phylogenetic diversity assuming no extinction (y-axes). Datapoints (N = 987 plots of 40 x 40 m) represent individual plant communities from the National Ecological Observatory Network. Colors indicate the % of species richness loss after dropping species that present evolutionary redundancy. SR = species richness; PD = phylogenetic diversity; PSR = phylogenetic species richness; PSV = phylogenetic species variability; MPD = mean phylogenetic pairwise distance; PSE = phylogenetic species evenness.
