## Supplemental Table 1 for "No branch left behind: tracking terrestrial biodiversity from a phylogenetic completeness perspective": Pinto-Ledezma_EtAl_SuppInfo_WebPanel_S1.docx

**JN Pinto-Ledezma *et al*. – Supporting Information**

**WebPanel 1. Description of the phylogenetic and geographical data**

We obtained phylogenetic hypotheses for terrestrial vertebrates from VertLife (<https://vertlife.org>), including amphibians (Jetz and Pyron 2018), squamates (Tonini *et al.* 2016), birds (Jetz *et al.* 2012), and mammals (Upham *et al.* 2019). Together these data comprise ~33,000 species, of which 7,238 are amphibians, 9,755 are squamates, 9,993 are birds, and 5,911 are mammals. Owing to the uncertainty in species relationships due to imputation of missing species, we obtained 10,000 trees from the Bayesian pseudo-posterior distribution for each vertebrate clade. Using these trees, we estimated the maximum clade credibility (MCC) tree for each clade using TreeAnnotator (Suchard *et al.* 2018). We used the MCC trees for further analyses. A phylogeny for seed plants was obtained from the recent published Spermatophyta mega-phylogeny (Smith and Brown 2018). This phylogeny comprises 353,185 species and is the most comprehensive phylogenetic tree for seed plants.

Geographical ranges of terrestrial vertebrate species were obtained from the IUCN (<https://www.iucnredlist.org/resources/spatial-data-download>) and BirdLife International (<http://datazone.birdlife.org/species/requestdis>). Geographical ranges of vascular plants were obtained from GreenMaps (Daru *et al.* 2021) that currently represent the most comprehensive database of geographical ranges for vascular plants globally. To obtain a species presence-absence matrix (PAM) we rasterized the species ranges into a grid of 1° × 1°degree—approximately 110 x 110 km near the equator—for each clade separately. The resulting PAMs were used for further analyzes and mapping purposes.

All data management and analyses were performed in R 4.1(R Core Team 2021) and using customized functions and the packages: ape (Paradis and Schliep 2019), phytools (Revell 2012), raster (Hijmans 2021), rgdal (Bivand *et al.* 2021), letsR (Vilela and Villalobos 2015), picante (Kembel *et al.* 2010), and tidyverse (Wickham *et al.* 2019). Bayesian MCP regressions were implemented in JAGS through the R package pcm (Lindeløv 2020). The R codes to implement the phylogenetic completeness approach and the Bayesian MCP regressions are available at GitHub (<https://github.com/jesusNPL/FITBITs>).

**WebReferences**

Bivand R, Keitt T, and Rowlingson B. 2021. rgdal: Bindings for the “Geospatial” Data Abstraction Library.

Daru BH, Davies TJ, Willis CG, *et al.* 2021. Widespread homogenization of plant communities in the Anthropocene. *Nat Commun* **12**: 6983.

Hijmans RJ. 2021. raster: Geographic Data Analysis and Modeling.

Jetz W and Pyron RA. 2018. The interplay of past diversification and evolutionary isolation with present imperilment across the amphibian tree of life. *Nat Ecol Evol* **2**: 850–8.

Jetz W, Thomas GH, Joy JB, *et al.* 2012. The global diversity of birds in space and time. *Nature* **491**: 444–8.

Kembel SW, Cowan PD, Helmus MR, *et al.* 2010. Picante: R tools for integrating phylogenies and ecology. *Bioinformatics* **26**: 1463–4.

Lindeløv JK. 2020. mcp: An R Package for Regression With Multiple Change Points. Open Science Framework.

Paradis E and Schliep K. 2019. ape 5.0: an environment for modern phylogenetics and evolutionary analyses in R. *Bioinformatics* **35**: 526–8.

R Core Team. 2021. R: A Language and Environment for Statistical Computing. Vienna, Austria: R Foundation for Statistical Computing.

Revell LJ. 2012. phytools: An R package for phylogenetic comparative biology (and other things). *Methods in Ecology and Evolution* **3**: 217–23.

Smith SA and Brown JW. 2018. Constructing a broadly inclusive seed plant phylogeny. *Am J Bot* **105**: 302–14.

Suchard MA, Lemey P, Baele G, *et al.* 2018. Bayesian phylogenetic and phylodynamic data integration using BEAST 1.10. *Virus Evolution* **4**.

Tonini JFR, Beard KH, Ferreira RB, *et al.* 2016. Fully-sampled phylogenies of squamates reveal evolutionary patterns in threat status. *Biological Conservation* **204**: 23–31.

Upham NS, Esselstyn JA, and Jetz W. 2019. Inferring the mammal tree: Species-level sets of phylogenies for questions in ecology, evolution, and conservation (AJ Tanentzap, Ed). *PLoS Biol* **17**: e3000494.

Vilela B, and Villalobos F. 2015. letsR: a new R package for data handling and analysis in macroecology. *Methods in Ecology and Evolution*.

Wickham H, Averick M, Bryan J, *et al.* 2019. Welcome to the tidyverse. *Journal of Open Source Software* **4**: 1686.
