## Supplemental Table 1 for "No branch left behind: tracking terrestrial biodiversity from a phylogenetic completeness perspective": Pinto-Ledezma_EtAl_SuppInfo_WebTable_S1.docx

**Pinto-Ledezma *et al*. – Supporting Information**

**WebTable 1.** Summary of the changing point analysis. Table shows the changing points on the terrestrial biodiversity identified by our Bayesian Multiple Changing Points (MCP) models. It also shows the observed diversity—in terms of number of species and phylogenetic diversity—and the associated diversity at each changing point. The lower credible intervals (CI [the 2.5% quantile of the posterior distribution]; boldfaced) were used as thresholds for further analyses. Note that the CP thresholds change depending on the clade. The observed number of species (second column) corresponds to the number of species sampled in each phylogenetic tree and might not represent each clade's true number of species.

| Clade | Observed  N species | Observed PD | Changing Point (CP) | Threshold (My) | CP  N species | CP  PD | %Spp  Loss | %PD  Loss |
| --- | --- | --- | --- | --- | --- | --- | --- | --- |
| Seed plants | 353185 | 4915453 | Mean | 2.65 | 289146 | 4841499 | 18.13 | 1.50 |
|  |  |  | **Lower-CI** | **1.76** | **312540** | **4887895** | **11.51** | **0.56** |
|  |  |  | Upper-CI | 3.559 | 268012 | 4778849 | 24.12 | 2.78 |
| Amphibians | 7238 | 131318 | Mean | 4.65 | 5833 | 127315 | 19.41 | 3.05 |
|  |  |  | **Lower-CI** | **4.43** | **5927** | **127736** | **18.11** | **2.73** |
|  |  |  | Upper-CI | 4.870 | 5744 | 126899 | 20.64 | 3.37 |
| Squamata | 9755 | 123103 | Mean CP | 3.07 | 7820 | 119807 | 19.84 | 2.68 |
|  |  |  | **Lower-CI** | **2.61** | **8141** | **120700** | **16.55** | **1.95** |
|  |  |  | Upper-CI | 3.525 | 7473 | 118686 | 23.39 | 3.59 |
| Birds | 9993 | 83101 | Mean | 0.90 | 9346 | 82754 | 6.47 | 0.42 |
|  |  |  | **Lower-CI** | **0.33** | **9859** | **83076** | **1.34** | **0.03** |
|  |  |  | Upper-CI | 1.502 | 8588 | 81843 | 14.06 | 1.51 |
| Mammals | 5911 | 35312 | Mean | 2.06 | 3739 | 33077 | 36.75 | 6.33 |
|  |  |  | **Lower-CI** | **0.31** | **5802** | **35289** | **1.84** | **0.06** |
|  |  |  | Upper-CI | 2.71 | 3237 | 32069 | 45.24 | 9.18 |
