## Supplemental Table 1 for "No branch left behind: tracking terrestrial biodiversity from a phylogenetic completeness perspective": Pinto-Ledezma_EtAl_SuppInfo_WebTable_S2.docx

**Pinto-Ledezma *et al*. – Supporting Information**

**WebTable 2.** Statistical summary of the comparison between conservation scenarios (phylogenetic completeness [PC] versus random loss [Rand]) at global and biome scale. For each comparison we computed the evidence ratio (*ER*) which represents the posterior probability of a hypothesis against an alternative hypothesis—hypothesis = PC > Rand. Values of *ER* greater than one suggest evidence for supporting the hypothesis that the PC scenario outperform the Rand scenario. Star indicate strong evidence in the favor the hypothesis. Hypothesis testing was performed using the in the probabilistic programming language Stan (Carpenter *et al.* 2017) through the R package brms (Bürkner 2017).

| **Biome** | **Clade** | **Estimate** | **Estimated**  **Error** | **CI**  **Lower** | **CI**  **Upper** | **Evidence**  **Ratio** | **Evidence** | **Posterior**  **Probability** | **Star** |
| --- | --- | --- | --- | --- | --- | --- | --- | --- | --- |
| Global | Seed plants | 3.348 | 0.070 | 3.234 | 3.459 | Inf | Strong | 1 | * |
| Global | Amphibians | 8.822 | 0.107 | 8.644 | 8.990 | Inf | Strong | 1 | * |
| Global | Squamata | 4.528 | 0.057 | 4.437 | 4.621 | Inf | Strong | 1 | * |
| Global | Birds | 1.090 | 0.007 | 1.079 | 1.102 | Inf | Strong | 1 | * |
| Global | Mammals | 0.051 | 0.007 | 0.039 | 0.062 | Inf | Strong | 1 | * |
| Boreal Forests/Taiga | Seed plants | -3.246 | 0.210 | -3.597 | -2.900 | 0.000 | NE | 0 |  |
| Boreal Forests/Taiga | Amphibians | 0.000 | 0.000 | 0.000 | 0.000 | 0.988 | Small | 0.497 |  |
| Boreal Forests/Taiga | Squamata | 0.000 | 0.000 | 0.000 | 0.000 | 0.984 | Small | 0.496 |  |
| Boreal Forests/Taiga | Birds | 0.608 | 0.020 | 0.575 | 0.640 | Inf | Strong | 1 | * |
| Boreal Forests/Taiga | Mammals | -0.398 | 0.077 | -0.528 | -0.267 | 0.000 | NE | 0 |  |
| Deserts & Xeric Shrublands | Seed plants | 3.443 | 0.161 | 3.177 | 3.714 | Inf | Strong | 1 | * |
| Deserts & Xeric Shrublands | Amphibians | 0.000 | 0.000 | 0.000 | 0.000 | 0.997 | Small | 0.49925 |  |
| Deserts & Xeric Shrublands | Squamata | 4.674 | 0.087 | 4.532 | 4.813 | Inf | Strong | 1 | * |
| Deserts & Xeric Shrublands | Birds | 1.368 | 0.020 | 1.336 | 1.400 | Inf | Strong | 1 | * |
| Deserts & Xeric Shrublands | Mammals | 0.000 | 0.000 | 0.000 | 0.000 | 1.012 | Small | 0.503 |  |
| Flooded Grasslands & Savannas | Seed plants | 5.345 | 0.986 | 3.484 | 6.650 | Inf | Strong | 1 | * |
| Flooded Grasslands & Savannas | Amphibians | 10.955 | 0.654 | 9.875 | 12.034 | Inf | Strong | 1 | * |
| Flooded Grasslands & Savannas | Squamata | 4.733 | 0.447 | 4.011 | 5.442 | Inf | Strong | 1 | * |
| Flooded Grasslands & Savannas | Birds | 1.030 | 0.039 | 0.967 | 1.093 | Inf | Strong | 1 | * |
| Flooded Grasslands & Savannas | Mammals | -0.036 | 0.105 | -0.199 | 0.140 | 0.538 | Small | 0.350 |  |
| Mangroves | Seed plants | 6.398 | 0.524 | 5.519 | 7.244 | Inf | Strong | 1 | * |
| Mangroves | Amphibians | 10.075 | 1.590 | 7.363 | 12.575 | Inf | Strong | 1 | * |
| Mangroves | Squamata | 3.581 | 0.859 | 2.178 | 5.008 | Inf | Strong | 1 | * |
| Mangroves | Birds | 0.920 | 0.067 | 0.811 | 1.029 | Inf | Strong | 1 | * |
| Mangroves | Mammals | 0.444 | 0.149 | 0.196 | 0.687 | 570.429 | Strong | 0.998 | * |
| Mediterranean Forests, Woodlands & Scrub | Seed plants | 4.855 | 0.196 | 4.533 | 5.178 | Inf | Strong | 1 | * |
| Mediterranean Forests, Woodlands & Scrub | Amphibians | 7.091 | 0.273 | 6.638 | 7.521 | Inf | Strong | 1 | * |
| Mediterranean Forests, Woodlands & Scrub | Squamata | 5.956 | 0.309 | 5.451 | 6.463 | Inf | Strong | 1 | * |
| Mediterranean Forests, Woodlands & Scrub | Birds | 0.752 | 0.047 | 0.674 | 0.831 | Inf | Strong | 1 | * |
| Mediterranean Forests, Woodlands & Scrub | Mammals | 0.000 | 0.000 | 0.000 | 0.000 | 1.027 | Small | 0.507 |  |
| Montane Grasslands & Shrublands | Seed plants | 3.617 | 0.247 | 3.206 | 4.019 | Inf | Strong | 1 | * |
| Montane Grasslands & Shrublands | Amphibians | 0.386 | 0.541 | -0.454 | 1.304 | 3.016 | Small | 0.751 |  |
| Montane Grasslands & Shrublands | Squamata | 6.460 | 0.332 | 5.901 | 7.001 | Inf | Strong | 1 | * |
| Montane Grasslands & Shrublands | Birds | 1.350 | 0.040 | 1.286 | 1.415 | Inf | Strong | 1 | * |
| Montane Grasslands & Shrublands | Mammals | 0.000 | 0.000 | 0.000 | 0.000 | 0.995 | Small | 0.499 |  |
| Temperate Broadleaf & Mixed Forests | Seed plants | -0.585 | 0.125 | -0.795 | -0.374 | 0.000 | NE | 0 |  |
| Temperate Broadleaf & Mixed Forests | Amphibians | 0.731 | 0.060 | 0.633 | 0.831 | Inf | Strong | 1 | * |
| Temperate Broadleaf & Mixed Forests | Squamata | 2.521 | 0.128 | 2.310 | 2.736 | Inf | Strong | 1 | * |
| Temperate Broadleaf & Mixed Forests | Birds | 1.412 | 0.015 | 1.388 | 1.437 | Inf | Strong | 1 | * |
| Temperate Broadleaf & Mixed Forests | Mammals | 0.071 | 0.010 | 0.055 | 0.089 | Inf | Strong | 1 | * |
| Temperate Conifer Forests | Seed plants | 0.086 | 0.190 | -0.222 | 0.405 | 2.042 | Small | 0.671 |  |
| Temperate Conifer Forests | Amphibians | 0.000 | 0.000 | 0.000 | 0.000 | 0.966 | Small | 0.49125 |  |
| Temperate Conifer Forests | Squamata | 7.149 | 0.183 | 6.840 | 7.442 | Inf | Strong | 1 | * |
| Temperate Conifer Forests | Birds | 0.608 | 0.037 | 0.548 | 0.668 | Inf | Strong | 1 | * |
| Temperate Conifer Forests | Mammals | -0.308 | 0.040 | -0.368 | -0.240 | 0.000 | NE | 0 |  |
| Temperate Grasslands, Savannas & Shrublands | Seed plants | -2.352 | 0.187 | -2.658 | -2.051 | 0.000 | NE | 0 |  |
| Temperate Grasslands, Savannas & Shrublands | Amphibians | 0.000 | 0.000 | 0.000 | 0.000 | 1.011 | Small | 0.50275 |  |
| Temperate Grasslands, Savannas & Shrublands | Squamata | 6.224 | 0.156 | 5.973 | 6.484 | Inf | Strong | 1 | * |
| Temperate Grasslands, Savannas & Shrublands | Birds | 1.088 | 0.020 | 1.055 | 1.120 | Inf | Strong | 1 | * |
| Temperate Grasslands, Savannas & Shrublands | Mammals | 0.056 | 0.006 | 0.045 | 0.067 | Inf | Strong | 1 | * |
| Tropical & Subtropical Coniferous Forests | Seed plants | 6.125 | 0.251 | 5.716 | 6.545 | Inf | Strong | 1 | * |
| Tropical & Subtropical Coniferous Forests | Amphibians | 7.253 | 0.917 | 5.739 | 8.777 | Inf | Strong | 1 | * |
| Tropical & Subtropical Coniferous Forests | Squamata | 5.770 | 0.632 | 4.765 | 6.804 | Inf | Strong | 1 | * |
| Tropical & Subtropical Coniferous Forests | Birds | 1.075 | 0.040 | 1.010 | 1.140 | Inf | Strong | 1 | * |
| Tropical & Subtropical Coniferous Forests | Mammals | 0.059 | 0.063 | -0.043 | 0.162 | 4.722 | Small | 0.825 |  |
| Tropical & Subtropical Dry Broadleaf Forests | Seed plants | 6.918 | 0.168 | 6.639 | 7.186 | Inf | Strong | 1 | * |
| Tropical & Subtropical Dry Broadleaf Forests | Amphibians | 11.152 | 0.264 | 10.723 | 11.579 | Inf | Strong | 1 | * |
| Tropical & Subtropical Dry Broadleaf Forests | Squamata | 3.608 | 0.219 | 3.249 | 3.969 | Inf | Strong | 1 | * |
| Tropical & Subtropical Dry Broadleaf Forests | Birds | 0.931 | 0.030 | 0.882 | 0.979 | Inf | Strong | 1 | * |
| Tropical & Subtropical Dry Broadleaf Forests | Mammals | -0.022 | 0.037 | -0.083 | 0.039 | 0.385 | Small | 0.278 |  |
| Tropical & Subtropical Grasslands, Savannas & Shrublands | Seed plants | 6.834 | 0.077 | 6.704 | 6.960 | Inf | Strong | 1 | * |
| Tropical & Subtropical Grasslands, Savannas & Shrublands | Amphibians | 10.201 | 0.203 | 9.862 | 10.539 | Inf | Strong | 1 | * |
| Tropical & Subtropical Grasslands, Savannas & Shrublands | Squamata | 6.430 | 0.104 | 6.259 | 6.598 | Inf | Strong | 1 | * |
| Tropical & Subtropical Grasslands, Savannas & Shrublands | Birds | 1.293 | 0.009 | 1.277 | 1.308 | Inf | Strong | 1 | * |
| Tropical & Subtropical Grasslands, Savannas & Shrublands | Mammals | 0.788 | 0.042 | 0.718 | 0.857 | Inf | Strong | 1 | * |
| Tropical & Subtropical Moist Broadleaf Forests | Seed plants | 8.169 | 0.061 | 8.069 | 8.269 | Inf | Strong | 1 | * |
| Tropical & Subtropical Moist Broadleaf Forests | Amphibians | 10.104 | 0.139 | 9.875 | 10.337 | Inf | Strong | 1 | * |
| Tropical & Subtropical Moist Broadleaf Forests | Squamata | 3.504 | 0.097 | 3.343 | 3.667 | Inf | Strong | 1 | * |
| Tropical & Subtropical Moist Broadleaf Forests | Birds | 1.085 | 0.011 | 1.067 | 1.102 | Inf | Strong | 1 | * |
| Tropical & Subtropical Moist Broadleaf Forests | Mammals | 0.184 | 0.018 | 0.155 | 0.212 | Inf | Strong | 1 | * |
| Tundra | Seed plants | -4.140 | 0.374 | -4.771 | -3.526 | 0.000 | NE | 0 |  |
| Tundra | Amphibians | 0.000 | 0.000 | 0.000 | 0.000 | 1.019 | Small | 0.50475 |  |
| Tundra | Squamata | 0.000 | 0.000 | 0.000 | 0.000 | 0.986 | Small | 0.4965 |  |
| Tundra | Birds | 0.941 | 0.034 | 0.887 | 0.997 | Inf | Strong | 1 | * |
| Tundra | Mammals | -1.588 | 0.163 | -1.858 | -1.323 | 0.000 | NE | 0 |  |

**WebReferences**

Bürkner P-C. 2017. **brms** : An *R* Package for Bayesian Multilevel Models Using *Stan*. *J Stat Soft* **80**.

Carpenter B, Gelman A, Hoffman MD, *et al.* 2017. *Stan* : A Probabilistic Programming Language. *J Stat Soft* **76**.
